# Post-translational regulation of the Numb/Notch pathway in neurogenesis and cancer by Dlk2

**DOI:** 10.1101/2023.07.20.549453

**Authors:** Stephanie.B Telerman, Russell.S Hamilton, Ben Shaw, Jordan.D Dimitrov, Ben Steventon, Anne.C Ferguson-Smith

## Abstract

**Perturbations in fundamental developmental pathways have a profound influence on tumorigenesis. Numb plays a pivotal role in vertebrate development, including neurogenesis and is a key negative regulator of Notch signaling**^1, 2^**. Perturbation of Numb expression affects brain morphology and cell fate**^3^**. While extensive research has been conducted on canonical Notch ligands, regulation by vertebrate-specific non-canonical ligands is not understood. Here we identify Delta like non-canonical Notch ligand 2/EGFL9 (Dlk2) as a regulator of zebrafish neurogenesis with mutants exhibiting early increase and subsequent depletion of neural stem cells, decreased radial glial cells density, impaired neuronal cell distribution, and hypersensitivity to stimuli mimicking the embryonic murine Numb/Numblike null phenotype**^3^**. Numb function is inactivated by aberrant phosphorylation**^4^**, and we show that Dlk2 protein exhibits a high affinity direct interaction with Numb, with loss of Dlk2 in zebrafish telencephalon increasing Numb Ser276 phosphorylation with a concomitant increase in Notch signaling. Patients with tumors exhibiting reduced levels of Dlk2 have a poorer prognosis, while overexpression of Dlk2 in human cancer cell lines reduces cell proliferation. Our findings identify Dlk2 as a key partner of Numb, a gatekeeper of its activity, and an important player in a network of protein interactions regulating both neurogenesis and cancer with potential therapeutic implications.**

## Introduction

Notch signaling is a highly conserved pathway involved in various developmental processes^5^. Several mechanisms have been reported in past decades, from canonical to non-canonical signaling, that can be ligand-dependent or independent^6^. Although the focus has mainly been on canonical Notch ligands, there is increasing evidence of the role of non-canonical ligands in the modulation of Notch function in vertebrates, particularly in the nervous system^7–9^,. In particular, a role for the vertebrate-specific non-canonical Notch ligand Delta-like homologue 1 (Dlk1) in the postnatal neurogenic niches in mice has indicated its active role in the maintenance of neural stem cells^10^. However, much less is known about the function and mechanism of its paralog Dlk2. Here we investigated the role of Dlk2 in neurogenesis and in the molecular mechanism that underlies its function. Using *in vivo* studies in zebrafish, cancer cell lines and biochemical assays, our work reveals a developmental pathway in which the atypical Notch ligand Dlk2 regulates vertebrate neurogenesis. Our findings show that Dlk2 directly interacts with the Notch antagonist Numb, abrogating its aberrant phosphorylation and enhancing its repression of Notch signaling.

## Results

### Dlk2 regulates zebrafish neurogenesis

Dlk2 encodes a unique transcript in the zebrafish genome. To evaluate the transcriptional landscape of Dlk2 through zebrafish brain development, publicly available single-cell RNA seq datasets derived from zebrafish heads harvested from embryonic to larval stages^11^ were analyzed (Fig. 1a, Supplementary Table 1). Dlk2 was detected in both mesoderm and ectoderm derived tissues (Fig. 1a, Supplementary Table 1). Within the brain, the strongest expression was located in the forebrain, specifically in the telencephalon, diencephalon, hypothalamus, and habenulae. Gene expression was found in neuronal sub-populations, including GABAergic and glutamatergic neurons, progenitors, and radial glial cells (Supplementary Table 1). Dlk2 is expressed in the eye (cornea) from 20 hours post-fertilization (hpf) and in the retina (RGC) from 2 days post-fertilization (dpf) (Fig. 1a, Supplementary Table 1). We corroborated these results by RT-qPCR, which show the steady increase of Dlk2 expression throughout embryonic development in zebrafish (Fig. 1b) and the transcripts in the telencephalon in 4 months post-fertilization (mpf) adults (Extended Fig. 1a). Immunostaining validated mRNA embryonic expression in the brain as well as in adult telencephalic neurons (Fig. 1c,d). Together, our data indicate that Dlk2 is expressed early in embryonic development and is maintained in the central nervous system in larval stage and in the adult telencephalon.

**Figure 1.**
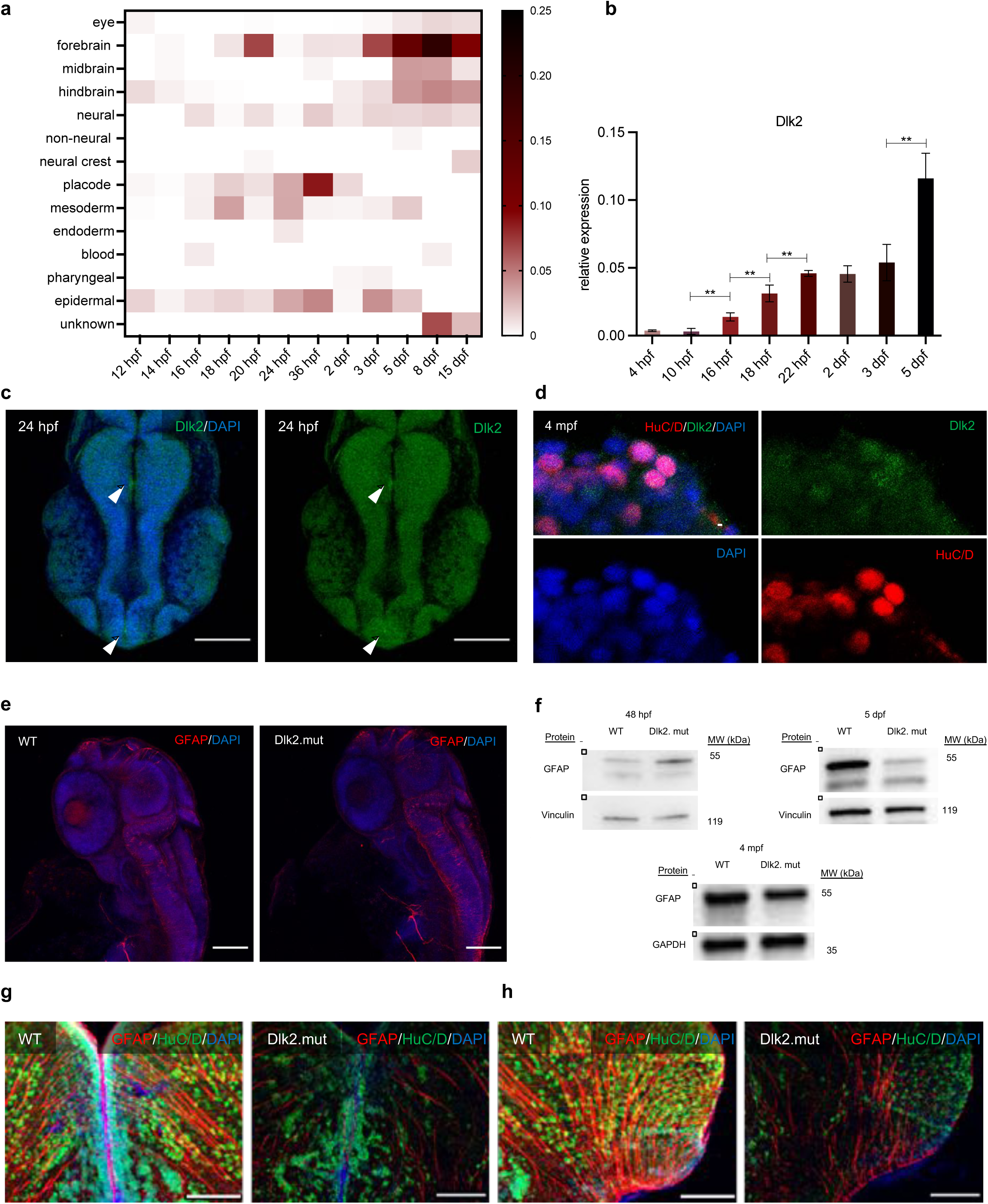
Dlk2 regulates neurogenesis. **a**, Heat map representing hierarchical clustering of differentially expressed Dlk2 gene in dissected zebrafish heads^11^ throughout development from 12 hpf to 15 dpf (single-cell sequencing datasets). **b**, RT–qPCR analysis of mRNA expression of *Dlk2* in the developing wild-type embryos. Data are mean of fold changes in relation to mobk13. Data are mean ± s.d (*n* ≥ 3 independent biological replicate experiments). **P ≤ 0.01 by unpaired Student’s *t*-test. **c**, Representative confocal images of wild-type zebrafish brain at 24 hpf (dorsal view) stained for Dlk2 (green, arrowheads indicating stronger Dlk2 expression) and DAPI (DNA, blue); (*n* ≥ 3 independent biological replicate experiments). Scale bars, 100 μm. **d**, Representative confocal images of wild-type zebrafish telencephalon pallium section at 4 mpf (coronal section) stained for Dlk2 (green), neuronal marker HuC/D (red) and DAPI (DNA, blue); (*n* ≥ 3 independent biological replicate experiments). Scale bar, 10 μm. **e**, Representative confocal images of wild-type (left) and Dlk2 mutant (right) zebrafish brain at 48 hpf (dorsal view) stained for GFAP (red) and DAPI (DNA, blue); (*n* ≥ 3 independent biological replicate experiments). Scale bars, 100 μm. **f**, Western blot of GFAP in zebrafish embryos at 48 hpf, 5 dpf, and 4 mpf in wild-type and Dlk2 mutant. Vinculin and GAPDH were used as loading controls. (*n* ≥ 3 independent biological replicate experiments). **g, h** Representative confocal images of wild-type zebrafish telencephalon: dorso-medial (**g**) and dorso-lateral (**h**) pallium section at 4 mpf (coronal section) stained for GFAP (red), neuronal marker HuC/D (green) and DAPI (DNA, blue). Scale bars, 100 μm. (*n* ≥ 3 independent biological replicate experiments).

To evaluate the role of Dlk2 in neurogenesis, we used a Dlk2 zebrafish mutant generated by the Sanger zebrafish mutant project, which lacks an essential splicing site before exon 4. We validated the Sanger sequencing, PCR genotyping, loss of transcript (RT-qPCR), and absence of protein (Extended Fig. 1b-f) in the mutant. The Dlk2 mutants were born at a normal Mendelian ratio and survived to adulthood with no effect on fertility or lifespan. The role of Dlk2 was investigated in secondary neurogenesis in the zebrafish mutants at 48 hpf by assessing the expression of the radial glial cells marker Glial Fibrillary acidic protein (GFAP). GFAP has previously been successfully used as a neural stem cell marker in zebrafish brain^12, 13^. We used this in combination with the pan-neuronal marker HuC/D (ELAVL3)^12, 13^. The mutants displayed increased GFAP expression at 48 hpf in the midbrain (Extended Fig. 1g) and hindbrain (Fig. 1e) confirmed by Western blot (Fig 1.f) as compared to wild-type zebrafish. This was followed by a reduction in protein levels measured at 5 dpf and adult forebrain (telencephalon) at 4 mpf, suggesting that Dlk2 is required for neurogenesis (Fig. 1f). The 4 mpf mutant telencephalon showed abnormal morphology, with a lower radial glial cell density (Fig. 1g,h, Extended Fig 2.a) and abnormal neuronal cell distribution with a clustering of the early-born neurons in the dorsal pallium and subpallium (Fig. 1g,h Extended Fig 2.b-g). As it is known that radial glial cells promote the generation of neuronal progenitors and play a fundamental role in their migration^14^, these data indicate that the loss of Dlk2 is likely to impair neural stem cell function with long-term consequences for adult telencephalon morphology.

**Figure 2.**
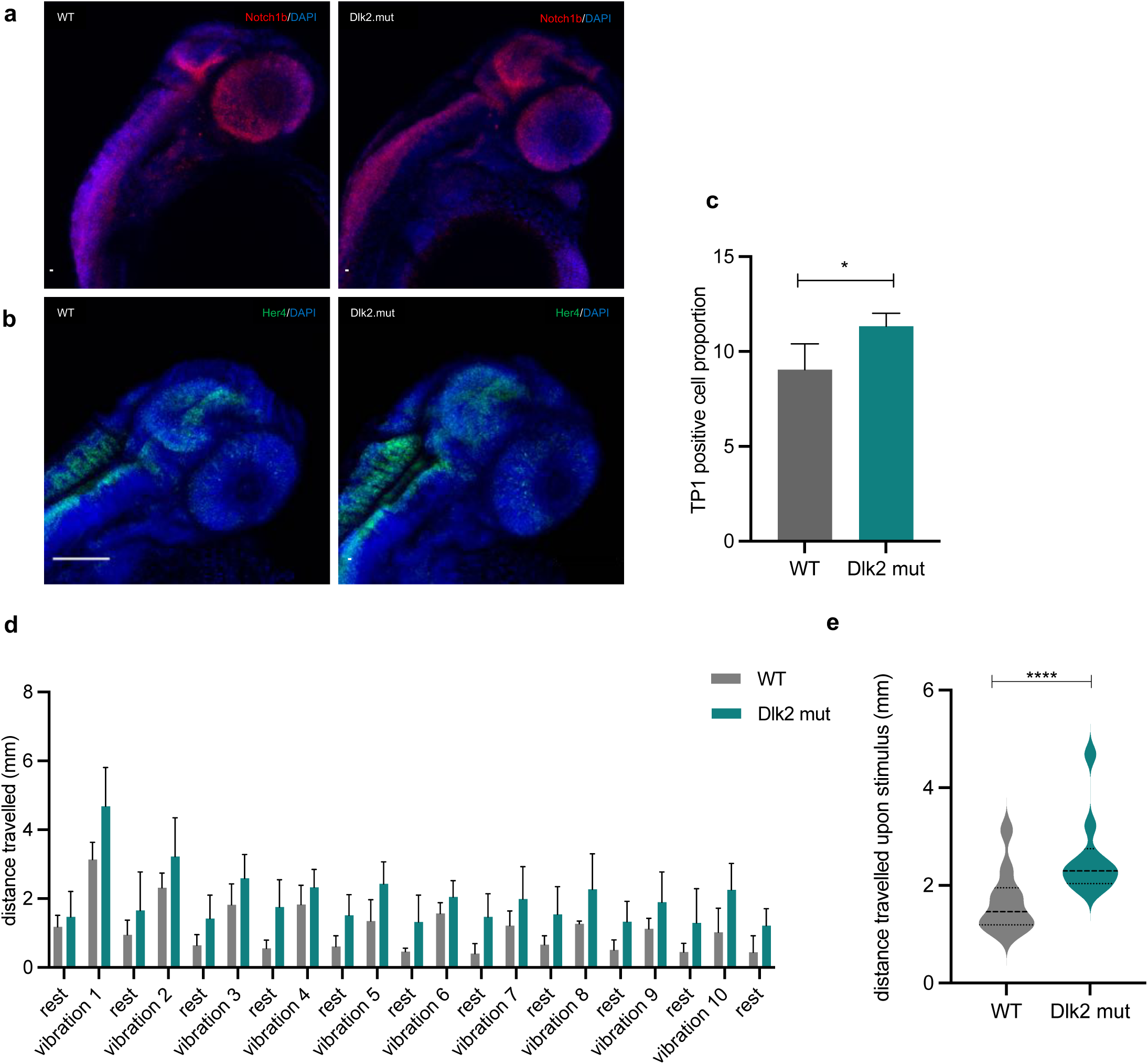
Dlk2 mutant display enhanced Notch signaling and hypersensitivity to stimuli. **a**, Representative confocal images of wild-type (left) and Dlk2 mutant (right) zebrafish brain at 48 hpf (dorsal view) labeled by *in situ* HCR for *Notch1b* (red) and DAPI (DNA, blue); (*n* ≥ 3 independent biological replicate experiments). Scale bars, 100 μm. **b,** Representative confocal images of wild-type (left) and Dlk2 mutant (right) zebrafish brain at 48 hpf (dorsal view) labeled by *in situ* HCR for *Her4* (green) and DAPI (DNA, blue); (*n* ≥ 3 independent biological replicate experiments). Scale bars, 100 μm**. c**, *Tp1:Venus* transgenic Notch reporter expression analyzed by flow cytometry in wild-type (left) and Dlk2 mutant (right) 26 hpf embryos); (*n* ≥ 3 independent biological replicate experiments); *P ≤ 0.05 by unpaired Student’s *t*-test. **d,** Representative startle experiment in wild-type (left) and Dlk2 mutant (right) zebrafish larvae at 7 dpf; (*n* ≥ 3 independent biological replicate experiments, each with n=12 zebrafish per genotype). **e**, Violin plot representative of the total distance traveled upon stimulus (excluding rest phases) measured in mm during the startle experiment (**d**) (*n* ≥ 3 independent biological replicate experiments, each with n=12 zebrafish per genotype). ****P ≤ 0.0001 by unpaired Student’s *t*-test.

### Dlk2 modulates Notch signaling and neuronal behavior

Dlk2 is required for secondary neurogenesis and the maintenance of radial glial cells, processes regulated by Notch signaling. We determined that there was a significant increase in the expression of relevant genes in the Dlk2-depleted zebrafish brain by assessing transcripts of Notch1b (Fig. 2a) and its target Her4 (Hes5 mammalian ortholog) (Fig. 2b). In addition, flow cytometry using the *Tp1:Venus* transgenic Notch reporter, which provides a readout for active Notch1 signaling^15^, on whole embryos at 26 hpf validated the increase in overall Notch signaling (Fig. 2c). Thus, the loss of Dlk2 results in increased levels of Notch receptor and downstream target transcripts.

To test if the neuropathological condition of the Dlk2 mutants affected behavior, changes in reaction to stimulus in larvae zebrafish at 7 dpf were quantified. Typically, zebrafish larvae exhibit a startle response upon exposure to tactile, acoustic, or visual stimuli and tend to avoid dark areas^16–18^. The startle response of zebrafish larvae was first examined in reaction to vibration stimuli (Fig. 2d,e). The Dlk2 mutant group had significantly elevated startle responses compared to wild-type zebrafish. The average total distance traveled was 2.57 ±0.81 mm for the mutants compared to 1.66 ±0.45 mm for controls. More precisely, the average distance traveled within 1 sec following the first startle stimulus was reduced to 3.13 ±0.51 mm compared to 4.68 ±1.12 mm for controls. A second behavioral test was performed, measuring the locomotion activity in light/dark transition cycles for the same larval stage. Indeed, it has been reported that larvae have a startle response with a brief period of elevated activity when exposed to a sudden change in light intensity^16^. Consistent with this, at the transition phase from the light to the dark phase, the activity of both the mutants and controls increased sharply (Extended Fig. 3a,b). The locomotion speed of the mutant group was significantly higher within 1 sec following the change of environment compared to controls. Altogether, these data suggest that neuronal and signaling alterations in Dlk2 mutants rendered the fish over-responsive to the vibration stimuli as well as to light intensity changes. These results were strikingly reminiscent of the Numb/Numblike knockout^3^ phenotype observed in cortical morphogenesis in mouse. Indeed, Numb is pivotal for the development of the mammalian brain, radial glial cell division and daughter cell fate specification^19–21^. This prompted us to explore the relationship between Dlk2 and Numb.

**Figure 3.**
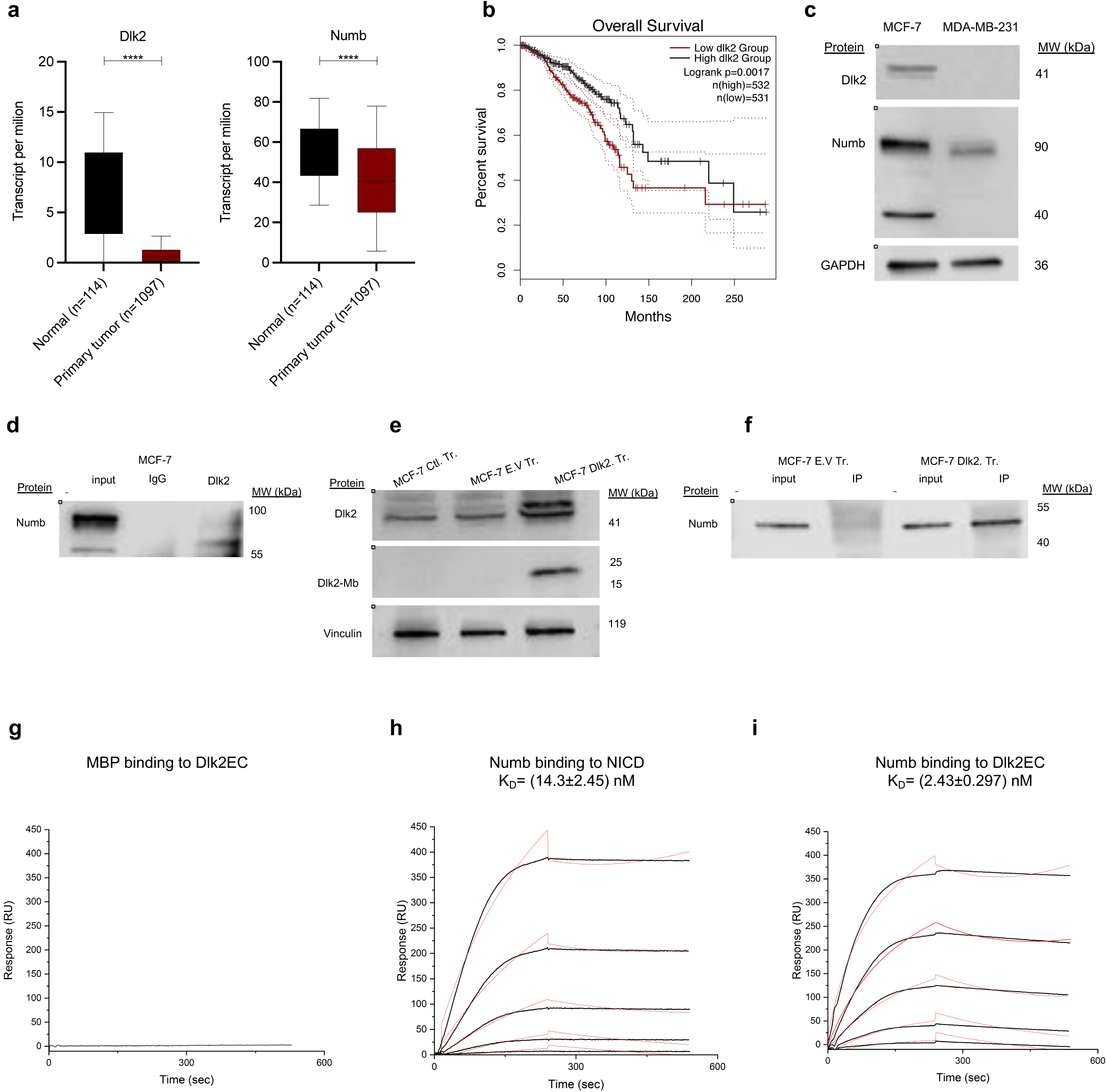
Dlk2 interacts directly with Numb. **a**, Expression level analysis of Dlk2 and Numb using UALCAN^44^ web-portal in normal breast tissue (n=114 patients) compared to invasive breast carcinoma (n=1097 patients) (P <1E-12 for Dlk2 as well as Numb). ****P ≤ 0.0001 by unpaired Student’s *t*-test. **b**, Kapan-Meier survival curves in invasive breast carcinoma on the basis of the UALCAN database^44^, comparing patients’ survival in subgroups high and low for Dlk2 expression. Survival analysis by Kapan-Meier curves and log-rank (Mantel-Cox) test P=0.0017. **c**, Western blot of Dlk2 and Numb in MCF-7 and MDA-MB-231 cell lines. GAPDH was used as a loading control. (*n* ≥ 3 independent biological replicate experiments). **d**, Co-immunoprecipitation (co-IP) experiment of Dlk2 in MCF-7 cell lysates. Equal amounts of extracts were immunoprecipitated either with Dlk2 antibody or IgG (control). Western blot for input along co-IP samples blotted with Numb antibody. (*n* ≥ 2 independent biological replicate experiments). **e**, Overexpression experiment of Dlk2 by transfection in MCF-7 cell line. Western blot of Dlk2, Dlk2-Mb (cleaved protein, corresponds to the molecular weight of the membrane-bound fragment) and Vinculin (loading control). Samples labeled as: transfection conditions control (Ctl. Tr), empty vector (E.V. Tr), and Dlk2 transfected cells (Dlk2. Tr). (*n* ≥ 3 independent biological replicate experiments). **f**, Co-immunoprecipitation (co-IP) experiment of Dlk2 in MCF-7 cell lysates comparing control (empty vector, E.V) transfected cells to Dlk2 transfected cells (Dlk2. Tr). Equal amounts of extracts were immunoprecipitated with DKK antibody against the DKK tag present in both the empty vector (control) and Dlk2 transfected cells (Dlk2. Tr). Western blot for input along co-IP samples blotted with Numb antibody. (*n* ≥ 2 independent biological replicate experiments). **g, h, i.,** Representative real-time interaction profiles obtained by surface plasmon resonance (SPR) kinetic analyses of the interaction between human Dlk2 extracellular domain (Dlk2EC) and Maltose-binding protein (MBP), negative control (**g**), human Notch1 intracellular domain (NICD) and Numb, positive control (**h**), and Dlk2 and Numb (**i**). The interaction profiles were obtained by binding increasing concentrations of MPB and Numb (0.019 to 20 nM) to DlkEC and NICD immobilized on sensor chips. The kinetic analyses were performed at 25 °C. Each graph represents experimental curves (black lines) and curves obtained by fitting the data using BIAevaluation software (red line). (*n* ≥ 2 independent biological replicate experiments).

### Dlk2 interacts with Numb

The Dlk2 mutant phenotype mimics a Numb/Numblike knockout^3^ and alters Notch signaling, both of which are known to play important roles in tumorigenesis. Consistent with this, Dlk2 was found to be differentially expressed in brain tumors of patients (Extended data Fig. 4a). Furthermore, in both low-grade gliomas and glioblastomas, Dlk2 was significantly decreased. Since Notch signaling is upregulated in these cancers, we specifically compared the expression of Dlk2 to Notch receptors and canonical ligands in glioblastoma (Extended data Fig. 4b). Strikingly, Dlk2 was significantly down-regulated. Jagged 2 was the only other significantly down-regulated gene. (Extended data Fig. 4b). Survival rates of patients with low (below the median value) and high expression levels of Dlk2 with low-grade glioma from the Cancer Genome Atlas (TGCA) database were compared, and we found that a lower expression correlates with poorer survival (p=0.001) (Extended data Fig. 4c).

**Figure 4.**
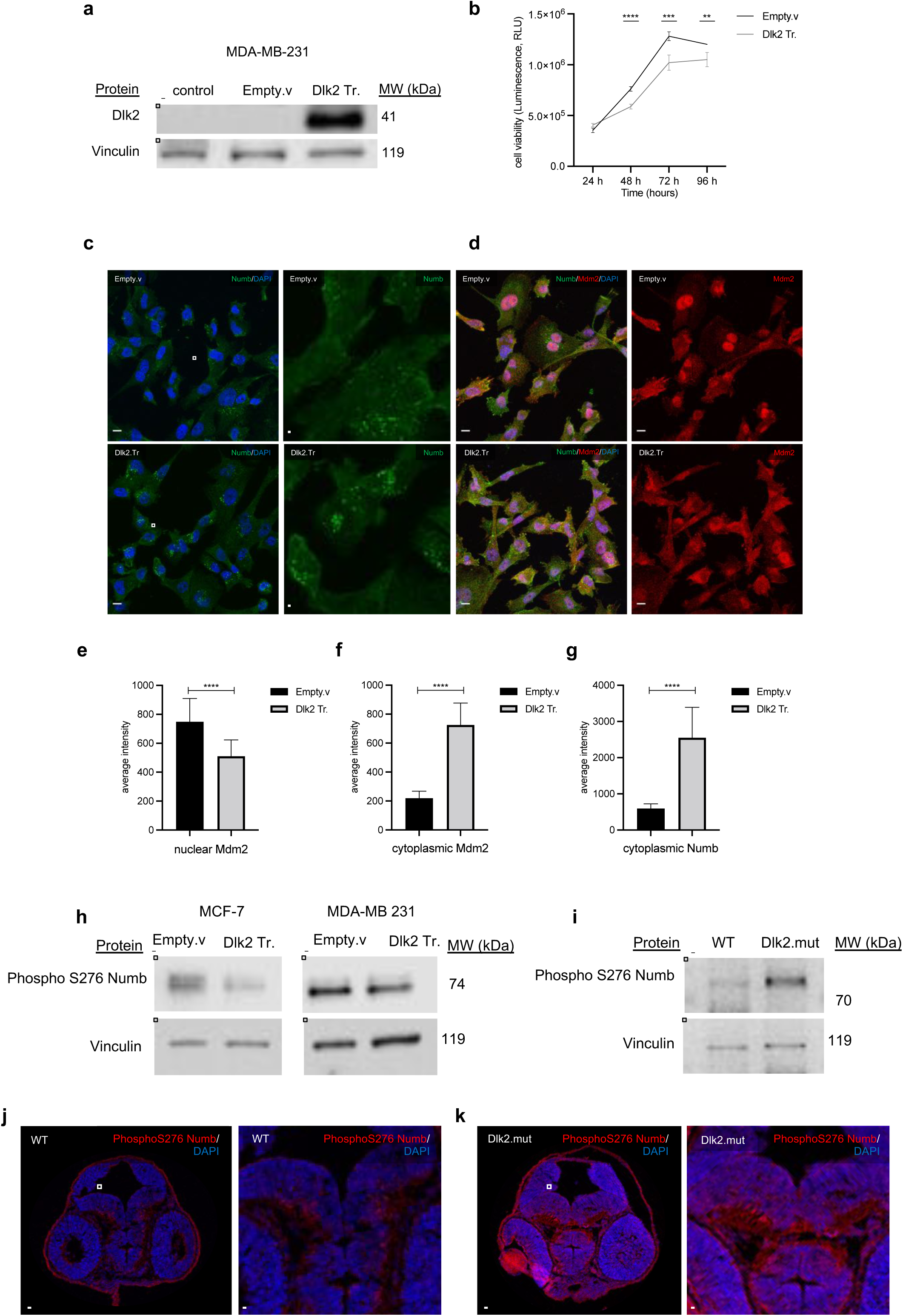
Dlk2 prevents Numb from aberrant Ser276 phosphorylation. **a**, Overexpression experiment of Dlk2 by transfection in MDA-MB-231cell line. Western blot of Dlk2, and Vinculin (loading control). For the samples: transfection conditions control, empty vector (Empty.v), and Dlk2 transfected cells (Dlk2. Tr). (*n* ≥ 2 independent biological replicate experiments). **b**, Cell viability assay of MDA-MB-231 cells comparing empty vector-transfected control cells (Empty.v) with Dlk2-transfected cells (Dlk2. Tr). Using the CellTiter-Glo assay, luminescence (RLU, relative light unit) was measured as an indicator of cell proliferation. Data are mean ± s.d taken every 24 h in replicate wells (*n* ≥ 3 independent biological replicate experiments). **c**, Representative confocal images of MDA-MB-231 cells comparing empty vector-transfected control cells (Empty.v) with Dlk2-transfected cells (Dlk2. Tr) stained for Numb (green) and DAPI (DNA, blue), left; Boxes outline enlarged area displayed on the right panels. Scale bars, 10 μm (*n* ≥ 3 independent biological replicate experiments). **d,** Representative confocal images of MDA-MB-231 cells comparing empty vector-transfected control cells (Empty.v) with Dlk2-transfected cells (Dlk2. Tr) stained for Numb (green), Mdm2 (red) and DAPI (DNA, blue). Scale bars, 10 μm (*n* ≥ 3 independent biological replicate experiments). **e,f,g,** Quantification of immunostainings representative of the confocal images displayed in (**d**). (*n*≥*114* cells for the empty vector control cells and *n*≥*130* for Dlk2-transfected cells). ****P ≤ 0.0001 by unpaired Student’s *t*-test. **h,** Western blot of phosphoSer276 Numb in MCF-7 (left) and MDA-MB-231 (right) cell lines. Vinculin (loading control). (*n* ≥ 3 independent biological replicate experiments). **i,** Western blot of phosphoSer276 Numb in adult zebrafish telencephalon comparing wild-type and Dlk2 mutant. Vinculin (loading control). (*n* ≥ 3 independent biological replicate experiments). **j,k** Representative confocal images of 48 hpf wild-type (**j**) and Dlk2 mutant (Dlk2.mut) (**k**) embryos (coronal cross section), stained for phosphoSer276 Numb (red) and DAPI (DNA, blue). Boxes outline the ventricle at the top of the thalamus in the midbrain. Scale bars, 100 μm (*n* ≥ 3 independent biological replicate experiments).

As Dlk2 is generally lowly expressed in these tumors, another tumor model was investigated to assess its potential interaction with Numb. Significant research on Numb in breast cancer has also shown it to be deregulated in these tumors, with a downstream effect on Notch signaling^22–24^. To gain initial insight into whether Dlk2 is likely to play a functional role in breast cancer progression in patients, publicly available datasets were queried. Analysis of gene expression data from TCGA showed that both Dlk2 and Numb expressions were significantly decreased in tumor tissue relative to normal breast tissue GTEx data (Fig. 3a). Assessment of expression levels relative to patient survival suggested that patients with a lower Dlk2 expression have a shortened survival compared to the high expression group (Fig. 3b).

Next, two breast cancer cell lines, MCF-7 and MDA-MB-231 were used to assess the expression and functional role of Dlk2. Dlk2 was expressed in MCF-7 cells but not in MDA-MB-231 (Fig. 3c). Numb protein was detected in both cell lines, though a slight reduction and different isoforms were noted in the latter (Fig. 3c). Co-immunoprecipitation of Dlk2 in the MCF-7 line detected a band corresponding to Numb (Fig. 3d). To confirm these results, we over-expressed Dlk2 by transfection into the MCF-7 line using a vector containing a DKK tag (Fig. 3e) and performed the co-IP against the DKK tag, which is also present in the empty vector control transfected MCF-7 line. A clear band was detected for Numb in the Dlk2-overexpressing cell line, indicating a potential interaction (Fig. 3f). Control co-IP in MDA-MB-231 cells which do not express Dlk2 confirmed the specificity of the interaction (Extended Fig. 4e). These data suggest that Dlk2 and Numb are part of the same complex.

In order to assess whether these proteins interact directly or indirectly, we gathered affinity and binding kinetics using surface plasmon resonance-based assay (SPR). We observed no specific interactions between Dlk2 and different negative controls (Fig. 3g; Extended data Fig. 5a-d). However, high-affinity interactions were observed between Dlk2 (Dlk2 extracellular domain (Dlk2EC)) and Numb with a KD value of ý2.4 nM, which was in a similar range to the interaction with the positive control Notch1 intracellular domain (NICD) (residues 2280-2550) and full-length Numb (KD ý14.3 nM) (Fig. 3h,i). The full-length Dlk2 bound to Numb with similar affinity to the Dlk2EC (data not included). A known target for Numb as a tumor suppressor is the E3 ligase Mdm2, which, upon interaction, is unable to bind p53^25^. It has also been reported that Mdm2 is able to target Numb for ubiquitination and subsequent proteasomal degradation. Accordingly, we hypothesized that Dlk2 might act to inhibit this interaction by potentially also binding to Mdm2. We used SPR technology to study the interaction between Dlk2 and Mdm2 (residues 119-438) (Extended data Fig. 5e). and observed a similar affinity to Dlk2 and Numb (KD ý3.5 nM). Moreover, further mapping experiments indicated that Dlk2 interacts with similar domains of Mdm2 (N-ter: residues 1-134 and central residues 134-334) as Numb, as we did not detect interaction with the C-terminal domain (residues 334-491) (Extended data Fig. 5f-h). Thus, Dlk2 is able to interact and form complexes with Numb and with Mdm2. However, despite validating the ubiquitination assay with an auto-ubiquitination of Mdm2 as well as Mdm2-mediated ubiquitination of p53 (Extended data Fig. 5i), as previously reported^26–29^, we could not confirm the degradation of Numb by Mdm2 (data not shown). Therefore, we hypothesize that Dlk2 functions in a different way within that complex.

### Dlk2 prevents Numb from aberrant phosphorylation

Since mutation of Dlk2 results in a Numb/Numblike phenotype, the biological relevance of the interaction between Dlk2 and Numb was assessed. Dlk2 was over-expressed by transfection into the MDA-MB-231 cell line (Fig. 4a). Analysis of the viability of these cells compared to the empty vector control line was measured daily as an indicator of cell proliferation, showing a significantly reduced capacity in the Dlk2-transfected cells (Fig. 4b). Reduced viability was also quantified in MCF-7 Dlk2-transfected cells (Extended Fig. 6a). This suggests a tumor suppressive role for the Dlk2 in this context. In order to consider the functional relationship with Numb, we tested whether increased Dlk2 expression could influence the activity of Numb protein in these cells. Immunostaining of Numb indicated a significantly increased expression in the cytoplasm of the Dlk2-transfected cells (Fig. 4c). Numb negatively regulates the function of Mdm2 and can shuttle the protein in and out of the nucleus^30^. Consistent with this, quantification of the expression levels of Mdm2 in the Dlk2-over-expressed transfected cells showed a decrease in Mdm2 in the nucleus with an accumulation of cytoplasmic expression (Fig. 4d-g), suggesting a regulatory role for Dlk2 in the interaction between Numb and Mdm2.

Numb was recently reported to be aberrantly phosphorylated on Ser 276 (S276) in breast cancer cells^4^. We confirmed these data in both MDA-MB-231 and MCF-7 (Fig. 4h). By contrast, in both lines over-expressing Dlk2, we detected a significant reduction of phosphorylated S276.

As shown previously^4^, Numb phosphorylation did not significantly affect the level of total Numb protein in these cells (Extended Fig. 6b). Thus, overexpression of Dlk2 is sufficient to diminish the post-translational modification of Numb and restore its activity. In addition, we tested the reverse condition in Dlk2 mutant zebrafish, by investigating the S276 Numb phosphorylation status, as this serine site is conserved between the two species (Extended Fig. 6c). Western blot analysis of adult zebrafish forebrain in Dlk2 mutants compared to controls showed a significantly increased level of phosphorylation (Fig. 4i) which is consistent with a negative regulatory role for Dlk2 in Numb S276 phosphorylation. Immunostaining using the same antibody confirmed these results in embryonic brains (Fig. 4j,k). Taken together, our results demonstrate a conserved biochemical function of Dlk2 in zebrafish and in mammals. Dlk2 interacts with Numb and Mdm2 and prevents Numb from aberrant phosphorylation, both in cancer cells and in neuronal development.

## Discussion

In this study, using zebrafish as a model to investigate neurogenesis, we have uncovered that the non-canonical Notch ligand Dlk2 regulates embryonic brain development. Compared to wild-type, Dlk2 mutant brains exhibited uncontrolled expansion of the radial glial cells, leading to improper radial glial morphology and density, along with impaired neuronal distribution in the adult forebrain. These phenotypes resemble those of the Numb/Numblike null neurogenesis phenotypes observed during embryonic cortical morphogenesis in mutant mice^3^. In addition to its role in neurogenesis Numb controls the stem cell compartment and asymmetric division in both vertebrates and Drosophila^19, 31, 32^. Hence, we assessed the potential regulation of Numb by Dlk2.

We found that Dlk2 resides in a complex with Numb and that it directly interacts with Numb with high affinity (KD;:2.4 nM). The cell-fate determinant activity of Numb relies highly on its interaction with various proteins. As an inhibitor of the Notch pathway, Numb binds to the active Notch intracellular domain (NICD) following internalization to prevent its translocation to the nucleus and transcription of downstream Notch targets^22, 33–35^. Through interaction with proteins of the endocytic machinery, Numb is also able to inhibit Notch1 activity by controlling post-endocytic sorting^1, 33, 36^. This negative regulation of Notch by Numb is crucial in the context of the central nervous system in which asymmetric localization in precursor cells generates two daughter cells with different fates (a Notch repressed cell as well as Numb negative cell)^36–38^. It is plausible that the interaction between Dlk2 and Numb potentiates its negative regulation of Notch in zebrafish neurogenesis to regulate the stem cell fate of radial glial cells.

Furthermore, in tumor biology, it was found that Numb regulates the tumor suppressor activity of p53, and is suggested to be a tumor suppressor itself by interacting with p53 and Mdm2^25, 26, 39, 40^. Alteration of Numb function contributes to human mammary carcinogenesis, and frequent loss of Numb expression has been found in breast tumors^22, 23^. Our findings show that Dlk2 interacts with both Mdm2 and Numb, leading us to explore further its role in tumor proliferation in breast cancer cells, as well as in cohorts of patients with breast cancer and low-grade glioma. In both cancer types, a decrease in Dlk2 expression was detected and linked to poorer survival prognosis. Thus, we concluded that Dlk2 is central in regulating Numb function in both neurogenesis and cancer. Because the protective role of Dlk2 is conserved from zebrafish to human breast cancer, we expect this targeted regulation to be of importance in settings beyond the scope of these models.

Phosphorylation of Numb has been reported to destabilize the NUMB-p53 interaction^41^ and, more specifically, aberrant phosphorylation on S276 residue renders Numb inactive in breast cancer, disrupting its tumor suppressor role^4^. This specific regulation of the phosphorylation of Numb was previously shown in other developmental systems, as constitutive S276 phosphorylation prevents Numb from recycling EGFR from the apical membrane of postnatal radial glial progenitors/stem cells in postnatal subependymal zone niche development^42, 43^. Here, we show that over-expression of Dlk2 in breast cancer cells led to a reduction in S276 phosphorylation of Numb. Conversely, the data in Dlk2 zebrafish mutants indicate an increase in S276 phosphorylation in the brain. The data uncover a previously unknown intracellular regulation of post-translational modifications of Numb by a vertebrate-specific non-canonical Notch ligand. These findings provide a novel mechanism of regulation of Numb activity and demonstrate that Dlk2 is necessary for determining homeostasis in both neurogenesis and tumor cell fate.

## Supporting information

Supplementary Table 1 _SB.Telerman

Supplementary Table 2 _SB.Telerman

## Materials and Methods

### Histology and Immunostaining

For immunostaining of zebrafish brain cryosections, tissue samples were fixed in 4% paraformaldehyde (PFA), equilibrated in 30% sucrose in PBS, embedded in optimal cutting temperature compound (OCT) (Agar Scientific), and stored at −80°C. Coronal cryosections (10 μm) were cut using a Microtome Leica Cryostat 3050S and stored at −80°C. For whole embryo tissue staining, and cell culture coverslips, samples were fixed in 4% PFA and stored at 4°C. For immunostaining of thick adult zebrafish brain sections, tissue samples were fixed in 4% PFA and stored in 100% methanol at −20°C. Samples were embedded in HistoGel (Fisher Scientific), and 70 μm thick sections were cut using a vibratome (VT1000S (L00-R21) and stored at 4°C. Samples were incubated in blocking buffer (0.1% Triton X-100, 0.1% Tween 20, 10% Fetal bovine serum (FBS)) for 1 hour at room temperature (RT) and incubated overnight at 4°C with primary antibodies in blocking buffer. Following repeated washing in PBS, samples were incubated for 1 hour at RT with Alexa Fluor-conjugated secondary antibodies in blocking buffer, washed, and mounted in Fluorescence Mounting Medium (Agilent Technologies). Primary antibodies were used at the indicated dilutions: Dlk2 (1:100, Abnova H00065989-B01P), GFAP (1:50, ZIRC ZRF-1), HuC/D (1:200, Abcam ab210554), DAPI (1:1000, Abcam ab228559), Numb (1:200, Abcam ab4147), PhosphoNumb Ser276 (1:200, Cell signaling Technology 4140S), Mdm2/HDM2 (1:200, R&D Systems MAB1244). Secondary antibodies were used at the indicated dilutions: AF488 Donkey anti-Goat (1:1000, Invitrogen A32814), AF555 Donkey anti-Rabbit (1:1000, Invitrogen A31572), AF647 Donkey anti-Mouse (1:1000, Invitrogen A31571). Microscopy was performed on a TCS SP8 Leica inverted confocal microscope at 10X, 20X or 40X magnification. Data analysis was performed using the image analysis softwares ImageJ and Imaris.

#### *In situ* hybridization

Embryos and larvae were fixed in 4% PFA overnight at 4°C. Next, samples were repeatedly washed in PBS-Tween-20 (PBST) before dehydration through a series of 25%, 50% and 75% methanol, stored in 100% methanol at −20°C, and processed for *in situ* RNA hybridization chain reaction (HCR). Samples were rehydrated by the methanol series, washed in PBST, and hybridized with 2 pmol HCR probes (Sigma-Aldrich) at 37°C overnight in a 30% formamide hybridization buffer. Next, repeated washing at 37°C was done with 30% formamide wash buffer. For probes detection, the annealing of fluorescent hairpins was done overnight at RT in amplification buffer. Next, repeated washing was done in 5X SSC 0.001 % Tween-20, and DAPI was added to the samples. Fluorescent hairpins and buffers were purchased from Molecular Technologies.

### Cell culture and transfection

MCF-7 and MDA-MB-231 cell lines were characterized and controlled for quality and maintained using standard tissue culture techniques. They were cultured in DMEM, high glucose (Gibco) supplemented with 10% FBS, 1% Penicillin-Streptomycin (Sigma-Aldrich); 0.2% insulin was added for MCF-7 cells. For transfection, cells were dissociated from 80% confluency the day before from a T-75 flask and seeded at 6.5 x 10^4^ cells to be at 70% confluency on the day of transfection on a 24-well plate. Lipofectamine 3000 reagents (Invitrogen) were used to transfect pCMV6-Entry Mammalian Expression control (PS100001, Origene) and Human Dlk2 pCMV6 (RC210622, Origene) vectors. Selection of transfected cells was performed using Neomycin. Cell viability of the transfected cells was measured by luminescence (RLU, relative light unit), quantifying the ATP production using the CellTiter-Glo Luminescent Cell Viability Assay (Promega) following the manufacturer’s instructions. Cells were seeded on a 96-well plate, and from 24 hours of culture, the luminescence assay was performed, with data acquisition done using the SpectraMax iD3 Plate Reader (Molecular Devices).

### Western blot

Cells were washed twice with PBS, collected with a cell scraper (Fisher Scientific) and lysed in ice-cold RIPA buffer (Sigma-Aldrich) supplemented with complete Mini EDTA-free Protease Inhibitor Cocktail tablets (Roche). Tissue samples were directly lysed in ice-cold supplemented RIPA buffer (Sigma-Aldrich). Samples were incubated on ice for 10 min, mechanically dissociated and incubated on ice for another 10 min. Following centrifugation at 4°C, supernatants were harvested on ice. Bradford (Bio-Rad) was used for protein quantitation and measured using a spectrophotometer. Laemmli Sample Buffer 4X (Bio-Rad) and β-mercaptoethanol (Sigma-Aldrich) 1/10^th^, were added to the samples before boiling at 95^◦^C. Protein extracts were separated on 4–20% polyacrylamide gels (Bio-Rad) and transferred to a PVDF membrane (Bio-Rad). Membranes were blocked for 1 hour at room temperature in 5% (w/v) non-fat milk (Marvel) or 5% (w/v) BSA (Sigma-Aldrich) depending on the antibody specifications, in 1× TBS and 0.1% Tween-20 (TBS-T) and incubated overnight at 4°C with the primary antibodies in blocking buffer. Membranes were washed three times for 20 min in TBS-T before incubation with horseradish peroxidase (HRP)-labeled secondary antibody in TBS-T at RT for 1 hour: Goat anti-Rabbit HRP (1:10,000, Agilent P0448), Rabbit anti-Goat HRP (1:10,000, Agilent P0449), Goat anti-Mouse HRP (1:10,000, Millipore AP308P). Following the washing of the membranes three times for 20 min in TBS-T, detection of antibodies was detected using the Amersham ECL Western Blotting Prime Blocking and Blotting Reagents (GE Healthcare).

### Immunoprecipitation

Cells were plated in T-75 flasks harvested at 80% confluency. Cells were then lysed in ice-cold lysis buffer (0.1% Triton X-100, 1% EDTA, PBS) for 10 min, mechanically dissociated and incubated on ice for another 10 min. Next, samples were centrifuged at 4°C for 20 min at 14.000 x g. Supernatants (2-3mg/ml) were harvested on ice with input samples, harvested (1/40^th^), Laemmli Sample Buffer (Bio-Rad) was added and boiled for 7 min at 95^◦^C. Supernatants were cleared by three consecutive incubation at 4°C with Protein A/G PLUS-Agarose beads (Santa Cruz). The elute was incubated overnight at 4°C with antibodies, and in parallel, beads were blocked overnight at 4 °C in blocking lysis buffer (BSA 0.5%, 0.1% Triton X-100, 1% EDTA, PBS). Next, beads were washed repeatedly with PBS and incubated at 4 °C with the elutes for 1h30. Beads were washed repeatedly with PBS, Laemmli Sample Buffer (Bio-Rad) added, and samples were boiled at 95^◦^C. Samples were centrifuged at 4°C, and supernatants were harvested and stored at −80^◦^C till further western blot analysis.

### RNA isolation and RT–qPCR

RNA was purified with the miRNeasy Micro Kit (Qiagen) according to the manufacturer’s instructions. Reverse transcription was done using SuperScript III Reverse Transcriptase (Thermo Fisher Scientific) with Random Primers following the manufacturer’s instructions. RT–qPCR reactions were performed using Brilliant III Ultra-Fast SYBR Green qPCR Master mix (Agilent Technologies) with the relevant primers (Sigma-Aldrich). Actbn2 (F-GGGCCTACCTCACTTTGAGC; R-CAACCAGTGCGGGAATTTCA), Mobk13 (F-AAGACTCCCAAAGAGTGCCC; R-CCCAGTTTGGCCACAGATGA), Dlk2 (F-GCCGATGTGTGACGACTGTG; R-GCTGTCTGGAGCACACGTAA). Data acquisition was done using the LightCycler480 machine (Roche) and QuantStudio 5 (Fisher Thermo Scientific) instruments. Analysis was performed by the delta-Ct method.

### PCR and Kompetitive Allele Specific PCR (KASP) genotyping assay

For rapid DNA extraction of zebrafish embryos/adult tail fin, samples were incubated in 25 μl of lysis buffer (25 mM NaOH with 0.2 mM EDTA) for 30 min at 95°C. Following rapid centrifugation, 25 μl of neutralization buffer (40mM Tris-HCl) was added to the samples, and after brief vortexing, DNA was used directly for PCR or genotyping assay or stored at −20°C. For PCR, a DNA loading buffer was added to the DNA samples. Agarose gel electrophoresis was done using gels made with TBE (Thermo Scientific), with DNA visualized under UV. Kompetitive Allele Specific PCR (KASP) (LGC group) genotyping assay was done using predesigned primers for the mutated sequence (A/T) of zebrafish Dlk2 (LGC group). Data acquisition was done using the LightCycler480 machine (Roche) and Quant Studio 5 (Fisher Thermo Scientific) instruments. Results were analyzed using the Endpoint Genotyping Software module of both instruments.

### *In vitro* ubiquitination assay

Reactions contained Tris 40 mM pH 7.4, 5mM Mg-ATP solution (Boston Biochem, B-20), 0.5 mM DTT, 0.6 μM ubiquitin (Boston Biochem, U-120), 225 nM E1-UBE-1 (Boston Biochem, E-305), 1.25 μM E2-UbcH5b (Boston Biochem, E3-622), and 2.5 μM p53 (Boston Biochem, SP-450). Reactions were started using E3 ligase MDM2/HDM2 400 ng (Boston Biochem, E3-204), incubated for 3 hours at 30°C in a water bath. The reactions were stopped by the addition of Laemmli Sample Buffer 4X (Bio-Rad) and β-mercaptoethanol (Sigma-Aldrich) 1/10^th^. Samples were analyzed using 4–20% polyacrylamide gels (Bio-Rad) followed by immunoblot with anti-Mdm2 (R&D, MAB1244) and anti-p53 antibodies (R&D, MAB1355).

### Zebrafish husbandry and behavioral assays

For Embryos were obtained from natural crosses and staged according to Kimmel et al^45^. Zebrafish lines were maintained under standard procedures in tanks with circulating system water at 28°C under 14:10 light: dark cycles. All zebrafish experiments were conducted in accordance with the University of Cambridge’s animal care guidelines under the Animals (Scientific Procedures) Act 1986 Amendment Regulations 2012, according to ethical review by the University of Cambridge Animal Welfare and Ethical Review Body (AWERB). All anesthesia was performed using Tricaine (MS-222, Sigma-Aldrich). Embryos were obtained from natural matings and grown at 28°C in E3 media at 28°C to the developmental stages of interest. The Dlk2 transgenic line was obtained from the zebrafish mutation project at the Sanger Institute, UK. Zebrafish mutants were outcrossed using wildtype zebrafish - Tüpfel long fin (TL) background and maintained. Tracking and recording, and data acquisition of larval (vibration startle and light/dark cycles assays) were done using the Zantiks MWP unit (Zantiks limited, Cambridge UK). The instrument was located in a quiet room with a controlled temperature of 27 ±1^◦^C. Each larva was transferred using a 2 ml pipette into wells of a clear 24-well plate (Thermo Fisher Scientific) containing 1 ml of E3 media per well. Fully detailed scripts of the experimental setups for both experiments are available via the link GitHub

### Flow cytometry

Embryos were harvested in E3 media and culled using Tricaine (MS-222, Sigma-Aldrich). Upon media change, deyolking buffer was added, and repeated pipetting followed by centrifugation was performed to remove the yolk. Disaggregation of zebrafish cells was done using the liberase enzyme (Roche). Incubation at room temperature for 20 min, including repeated pipetting, was concluded by the addition of fetal bovine serum (FBS) to the samples. Following centrifugation, cells were filtered through a 30-mm cell strainer, resuspended in ice-cold PBS, and labeled with DAPI to exclude dead cells. Data acquisition was performed using the Attune Flow cytometer (Thermo Fisher Scientific). For gate setting and compensation, unlabeled, single-labeled cells were used as controls. Data analysis was performed using FlowJo software, version 10 (Tree Star, Ashland, OR).

### Biacore

Kinetics of Human Dlk2 full length, Dlk2 extracellular domain (Dlk2EC), and Notch1 intracellular domain (NICD) binding to Human Mdm2 and Numb was evaluated by surface plasmon resonance-(SPR) based assay (Biacore 2000, Biacore Cytiva, Uppsala, Sweden). Dlk2 full length (Abnova, H00065989-P01), Dlk2EC (Abcam, ab109347), and NICD (Abcam, ab276343) were immobilized on a CM5 sensor chip using an amino-coupling kit as recommended by the manufacturer (Biacore, Cytiva), and a control flowcell was mock-immobilized. The experiments were performed using a running buffer made of 150 mM NaCl and 10 mM Hepes (pH 7.4), filtered through 0.22 µm filter and degassed under vacuum. The following proteins were injected to evaluate the binding kinetics of the interactions with Dlk2 full length, Dlk2EC, and NICD: Mdm2 (MDM2 protein Gln119-Leu438 Origene TP761916; MDM2 protein-full length Abcam, ab82080; Mdm2 truncated N-ter, central, C-ter domains (gift of A. Lespagnol), Numb full-length (Origene, TP761513), Human B7-H2-Fc Chimera (Biolegend, 599904), Human SNAP43 (SNAPC1) (Origene, TP760339), Human MBP (gift of A. Lespagnol), and Human MTERF (MTERF1) (Origene, TP761846). Proteins injected were serially diluted (two-fold each step) in running buffer to concentrations ranging from 200 to 0.019 nM prior to injection and introduced to the flowcells at a flow rate of 10 μl/min.

Regeneration of the chip surface was achieved by brief exposure (30 s) to a solution containing 3 M solution of KSCN (Sigma-Aldrich). BIAevaluation software (version 4.1.1, Biacore) was used to evaluate the kinetic rate constants, applying Langmuir binding model global kinetics analyses.

### In silico analysis using single-cell sequencing zebrafish datasets

Reanalysis of the single-cell RNA-Seq data from Raj et al, 2020^11^ was performed using R (v4.2.1) and R-Studio (v2022.12.+353). From the Gene Expression Omnibus (GEO) data accession number GSE158142; the Robject files were downloaded for each time point and loaded into Seurat^46^ (v4.3.0). The sample sheet containing metadata (RDS, Stage, Units, ClusteringResolution) for the Robjects was also downloaded in order to match the published annotations and resolutions for the dimensionality reduction. UMAP plots at the published resolution were plotted for specific genes of interest using FeaturePlot(). Average expression for published clusters and resolutions were extracted using AverageExpression() and written to XLS format files.

### In silico analysis using TCGA patients datasets

RNA expression levels and survival data were downloaded from the UALCAN web-portal^44^ as well as http://gepia2.cancer-pku.cn and www.oncolnc.org web-portals to assess the Cancer Genome Atlas/Genotype-Tissue Expression (TCGA/GTEx) database. Data with the accession number GSE4290 were analyzed from the TCGA database https://portal. gdc.cancer.gov.

### Code availability

Analysis of the single-cell RNA-Seq data^11^ accession number GSE158142 is available via GitHub https://github.com/darogan/AFS_st547_0002. Results were based on data downloaded from TCGA, accession number GSE4290 (https://portal. gdc.cancer.gov). We have provided shell scripts, and a Rscript download and recreated the gene-specific analyses via GitHub; We also provide gene-specific data and plot files.

### Quantification and statistical analysis

Statistical analyses were performed using GraphPad-Prism software. Unless stated otherwise, statistical significance was determined by the unpaired Student’s t-test to quantify differences between experimental groups. (ns, not significant, *P ≤ 0.05, **P ≤ 0.01, ***P ≤ 0.001, ****P ≤ 0.0001).

## Acknowledgments

We thank A. Flemming, A. Ashcroft, M. Keegan for early discussions. S. Patel, A. D. Hay, and all the members of the Ferguson-Smith laboratory for helpful discussions and for excellent suggestions; B. Buddenberg and J. Dagget from Zantiks Ltd, for their support with the zebrafish behavior tracking experiments. Biological Services Unit staff for assistance and animal maintenance in the zebrafish facility (University of Cambridge, PDN). This work was funded by the UK Medical Research Council.

## Author contributions

S.B.T. and A.C.F-S. conceived the study and designed the experiments. S.B.T, performed the *in vitro* and *in vivo* experiments, with the support of J.D.D for the SPR. R.S.H performed computational analysis of zebrafish single-cell RNA seq data and supervised analysis of patient data. B.S contributed to the initial phase of the zebrafish study. B.S contributed to the design of zebrafish study. S.B.T. and B.S and A.C.F-S. wrote the manuscript, with contributions from all authors.

## Competing interests

The authors declare no competing financial interests.

**Supplementary Table 1.** Single-cell RNAseq zebrafish datasets for Dlk2 expression

**Supplementary Table 2.** *in situ* HCR primers

## Extended data figure legends

**Extended Figure 1.**
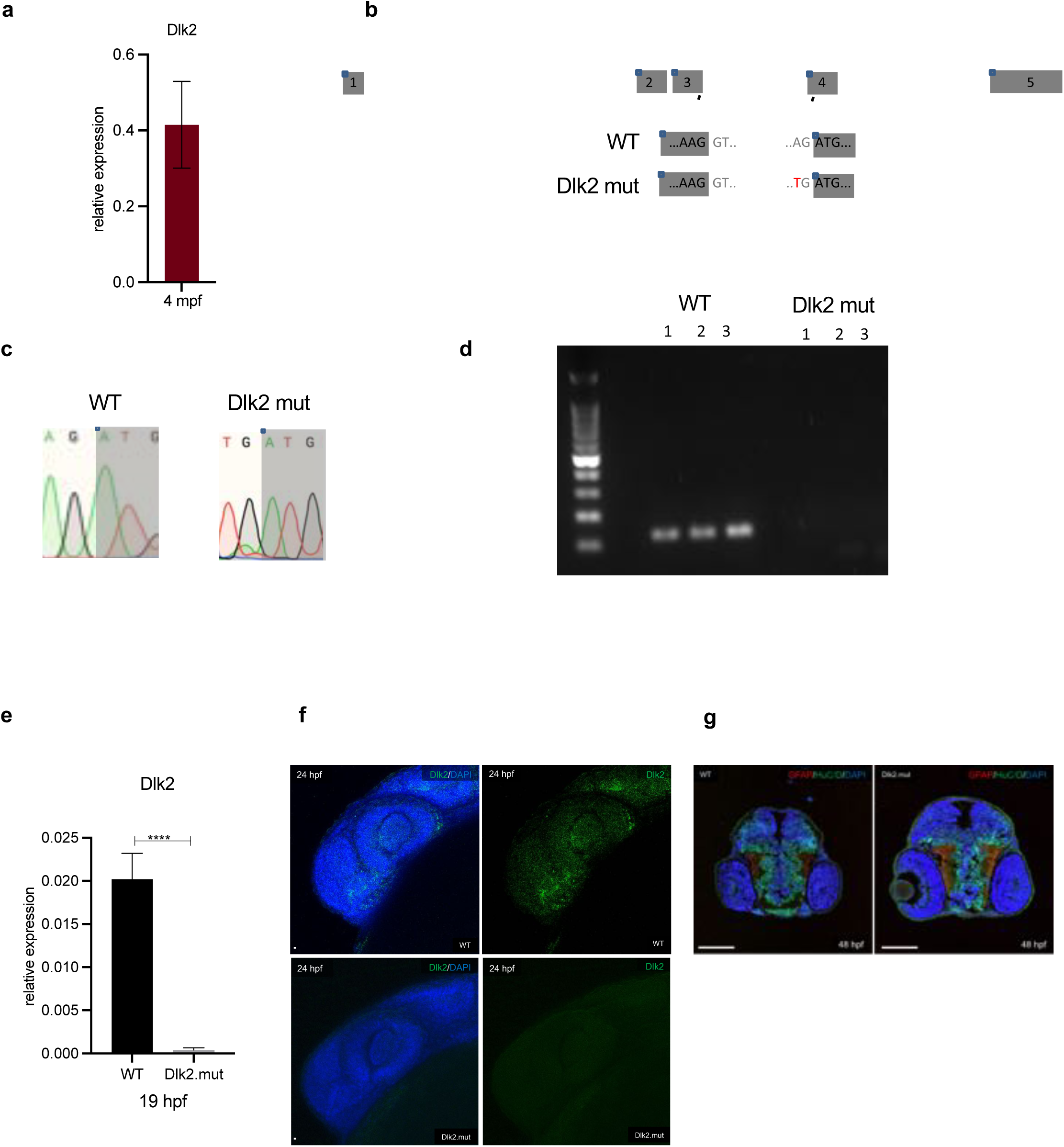
Characterization of Dlk2 mutant. **a**, RT–qPCR analysis of wild-type mRNA expression of *Dlk2* in the adult (4 mpf) telencephalon. Data are mean of fold changes in relation to actb2. Data are mean ± s.d (*n* ≥ 3 independent biological replicate experiments). **b**, Diagram of *Dlk2* zebrafish DNA sequence. Grey boxes represent exons; essential splice site sequences between exon 3 and 4 are shown, with the mutation A>T present in Dlk2 mutant. **c**, Sanger sequencing chromatograms of the essential splice site intronic sequences before exon 4 of Dlk2 comparing the wild-type with the Dlk2 mutant. **d**, Genotyping PCR comparing the wild-type with the Dlk2 mutant. **e**, RT–qPCR analysis of mRNA expression of *Dlk2* in the embryo (19 hpf) in wild-type and Dlk2 mutant. Data are mean of fold changes in relation to mobfk13. Data are mean ± s.d (*n* ≥ 3 independent biological replicate experiments). ****P ≤ 0.0001 by unpaired Student’s *t*-test. **f**, Representative confocal images of 24 hpf wild-type and Dlk2 mutant embryos (wholemount), stained for Dlk2 (green) and DAPI (DNA, blue), (*n* ≥ 3 independent biological replicate experiments). **g**, Representative confocal images of 48 hpf wild-type and Dlk2 mutant embryos (coronal cross section), stained for HuC/D (green), GFAP (red) and DAPI (DNA, blue), (*n* ≥ 3 independent biological replicate experiments).

**Extended Figure 2.**
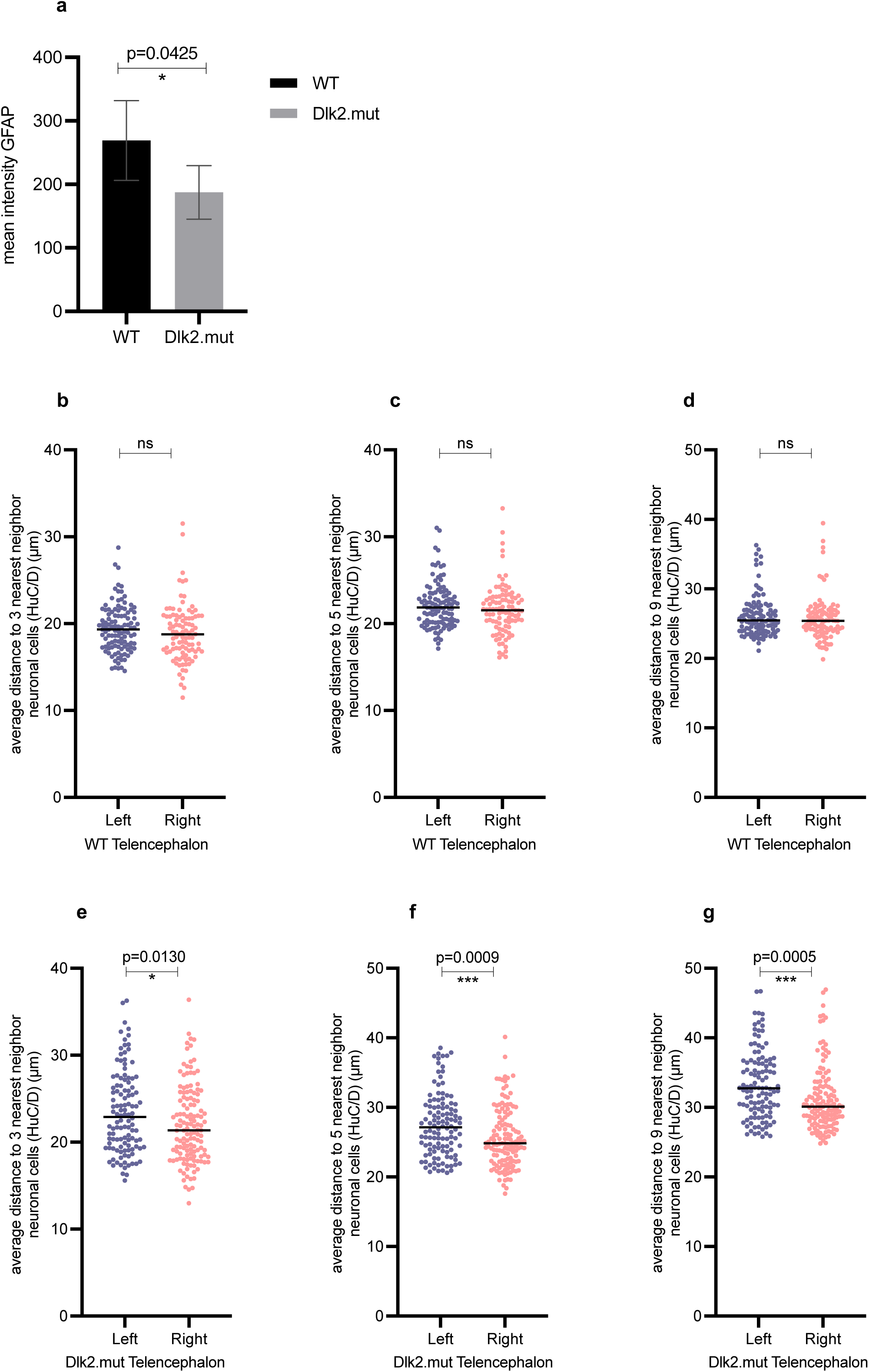
Altered neuronal populations in the telencephalon of adult Dlk2 mutants. **a,** Fiji analysis of GFAP intensity mean in adult telencephalon in wild-type and Dlk2 mutant zebrafish at 4 mpf (*n* ≥ 4 independent biological replicate experiments). *P = 0.0425 by unpaired Student’s *t*-test. **b-g** Plot representative of the average distance (μm) to the 3 (**a,e**), 5 (**c,f**), 9 (**d,g**) nearest neighbor neuronal cells (HuC/D used as a marker) in wild-type and Dlk2 mutant zebrafish at 4 mpf telencephalon (subpallium). Distribution is compared between the left and right hemispheres as an assessment of symmetry, (*n* ≥ 4 independent biological replicate experiments). *P = 0.0130 for 3 nearest neighbor cells, ***P = 0.0009 for 5 nearest neighbor cells, ***P = 0.0005 for 9 nearest neighbor cells, by unpaired Student’s *t*-test.

**Extended Figure 3.**
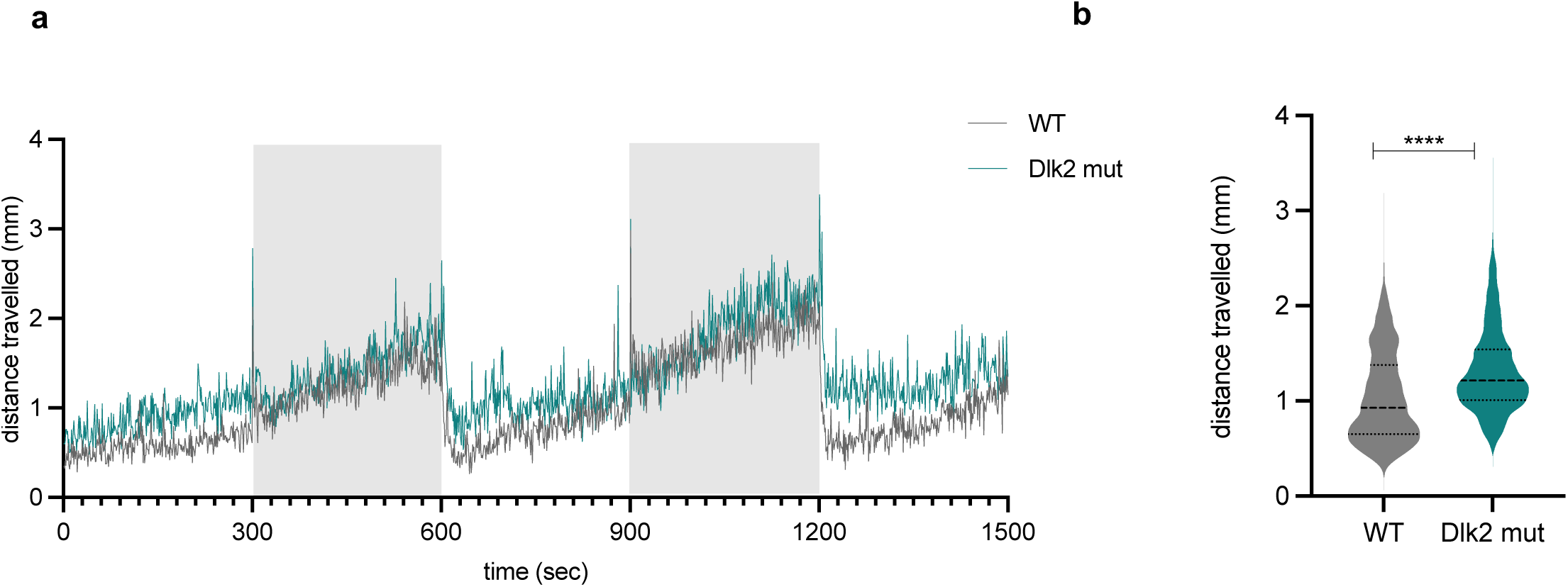
Dlk2 mutants are hypersensitive to light changes. **a,** Representative startle experiment upon light changes through time (grey boxes: light off) in wild-type (left) and Dlk2 mutant (right) zebrafish larvae at 7 dpf; (*n* ≥ 3 independent biological replicate experiments, each with n=12 zebrafish per genotype). **b,** Violin plot representative of the total distance traveled measured in mm during the experiment (**a**) (*n* ≥ 3 independent biological replicate experiments, each with n=12 zebrafish per genotype). ****P ≤ 0.0001 by unpaired Student’s *t*-test

**Extended Figure 4.**
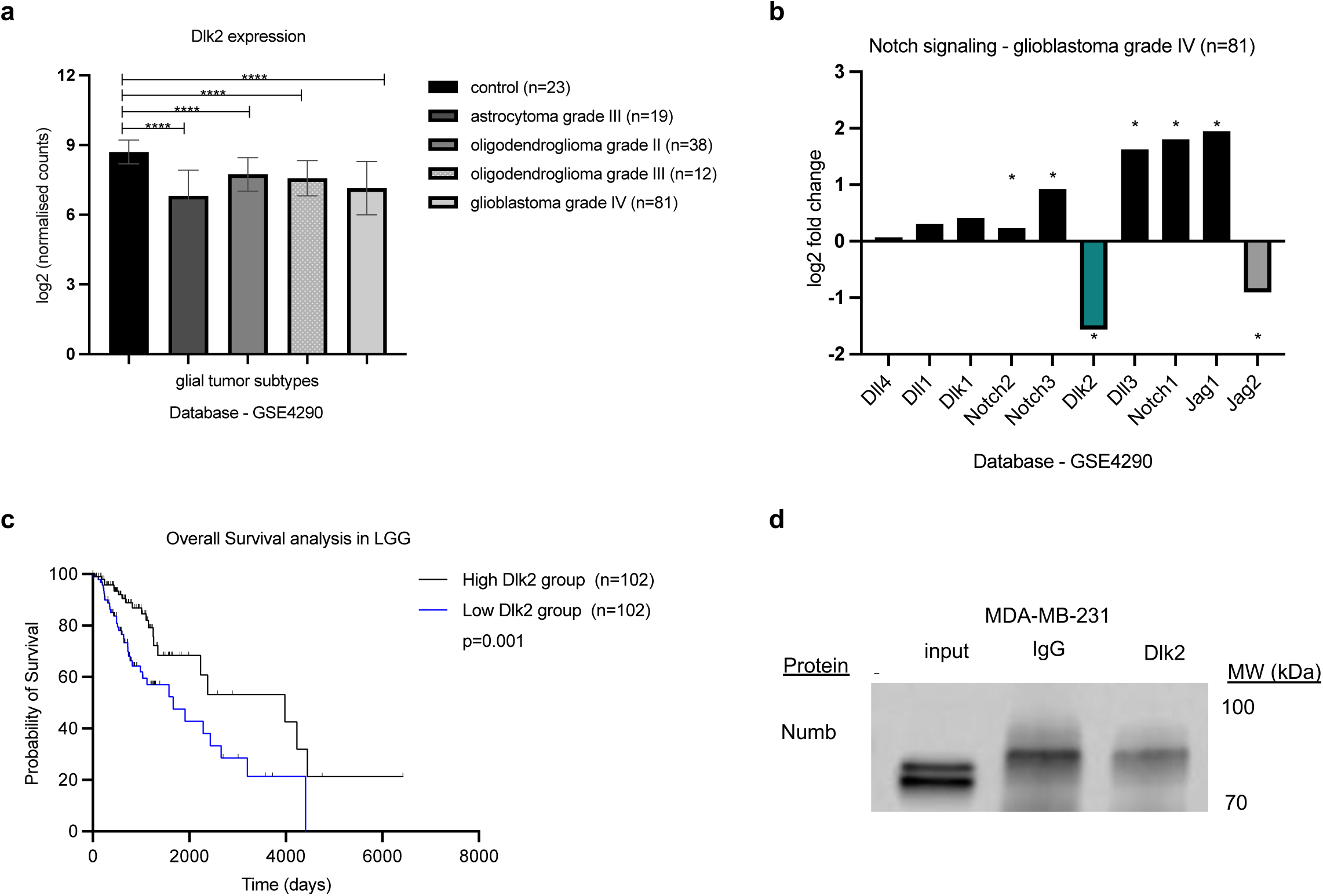
Dlk2 expression is downregulated in gliomas. **a**, Expression level analysis of Dlk2 and Numb using TCGA and GTEX datasets (Gepia2 database) in normal brain tissue (n=30) compared to astrocytoma grade III (n=19), oligodendroglioma grade II (n=38), oligodendroglioma grade III (n=12), and glioblastoma grade IV (n=81). ****P ≤ 0.0001 by unpaired Student’s *t*-test**. b**, Fold change expression of notch signaling (receptors and ligands) in glioblastoma grade IV (n=81) TCGA datasets displayed on (**a**). **c,** Survival analysis by Kapan-Meier curves and log-rank (Mantel-Cox) test in low-grade glioma on the basis of the TCGA database http://www.oncolnc.org, comparing patients’ survival in subgroups high and low for Dlk2 expression, **P = 0.001. **d,** Co-immunoprecipitation (co-IP) experiment MDA-MB-231 cell lysates that do not express Dlk2. This experiment is in comparison to the co-IP performed in MCF-7 lines that express Dlk2 (Figure 3e), to be used as a negative control. Equal amounts of extracts were immunoprecipitated either with Dlk2 antibody or IgG (control). Western blot for input along co-IP samples blotted with Numb antibody. No specific Numb band was detected in the Dlk2 IP compared to the control. (*n* ≥ 2 independent biological replicate experiments).

**Extended Figure 5.**
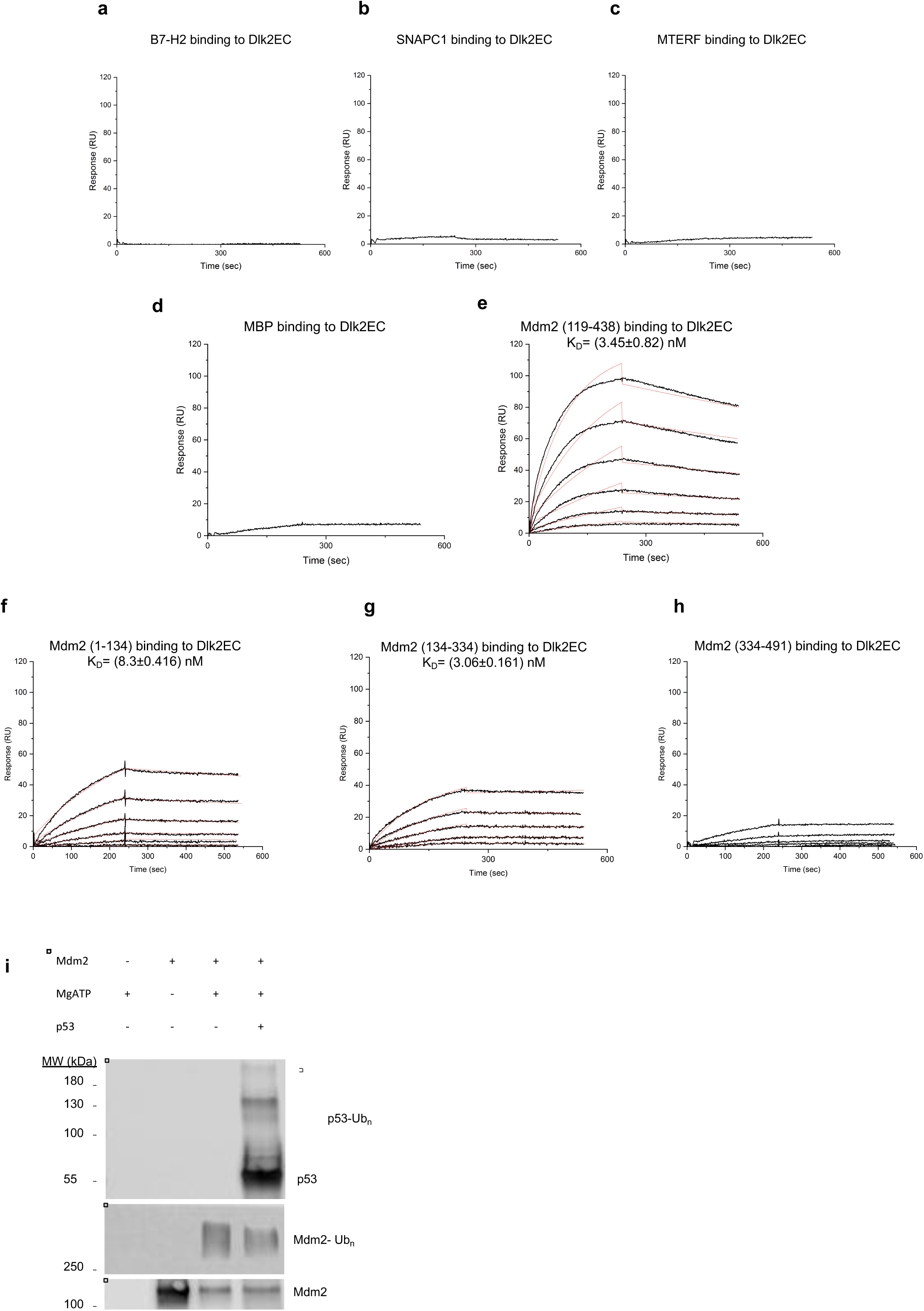
Dlk2 interacts directly with Mdm2. **a,b,c,** Representative real-time interaction profiles obtained by surface plasmon resonance (SPR) kinetic analyses of the interaction between human Dlk2 extracellular domain (Dlk2EC) and different negative controls: Human B7H2 (**a**), Human SNAPC1 (**b**), Human MTERF (**c**). The interaction profiles were obtained by binding increasing concentrations of B7H2, SNAPC1 and MTERF (0.019 to 20 nM) to DlkEC immobilized on sensor chips. The kinetic analyses were performed at 25 °C. Each graph represents experimental curves (black lines); as no interaction was detected, no fitting of the data is presented. **d,e,** Representative real-time interaction profiles obtained by surface plasmon resonance (SPR) kinetic analyses of the interaction between human Dlk2 extracellular domain (Dlk2EC, residues 1-306) and the negative control Human MBP (**d**) and Human Mdm2 (residues 119-438) (**e**). The interaction profiles were obtained by binding increasing concentrations of MBP and Mdm2 (0.19 to 200 nM) to DlkEC immobilized on sensor chips. The kinetic analyses were performed at 25 °C. Each graph represents experimental curves (black lines) and curves obtained by fitting the data using BIAevaluation software (red line). (*n* ≥ 2 independent biological replicate experiments). **f,g,h,** Representative real-time interaction profiles obtained by surface plasmon resonance (SPR) kinetic analyses of the interaction between human Dlk2 extracellular domain (Dlk2EC) and Human Mdm2 N-ter (residues 1-134) (**f**), Human Mdm2 central domain (residues 134-334) (**g**), and Human Mdm2 C-ter (residues 334-491) (**h**). The interaction profiles were obtained by binding increasing concentrations of Human Mdm2 N-ter, Human Mdm2 central domain, Human Mdm2 C-ter (0.19 to 200 nM) to DlkEC immobilized on sensor chips. The kinetic analyses were performed at 25 °C. Each graph represents experimental curves (black lines) and curves obtained by fitting the data using BIAevaluation software (red line). (*n* ≥ 2 independent biological replicate experiments). **i,** *in vitro* ubiquitination assay, p53 was incubated with ubiquitin, E1, E2 (UbcH5b), and the presence or absence of Mg-ATP and Mdm2 as indicated. Ubiquitination was detected by Western blot using anti-Mdm2 antibody and anti-p53 antibody (*n* ≥ 2 independent biological replicate experiments).

**Extended Figure 6.**
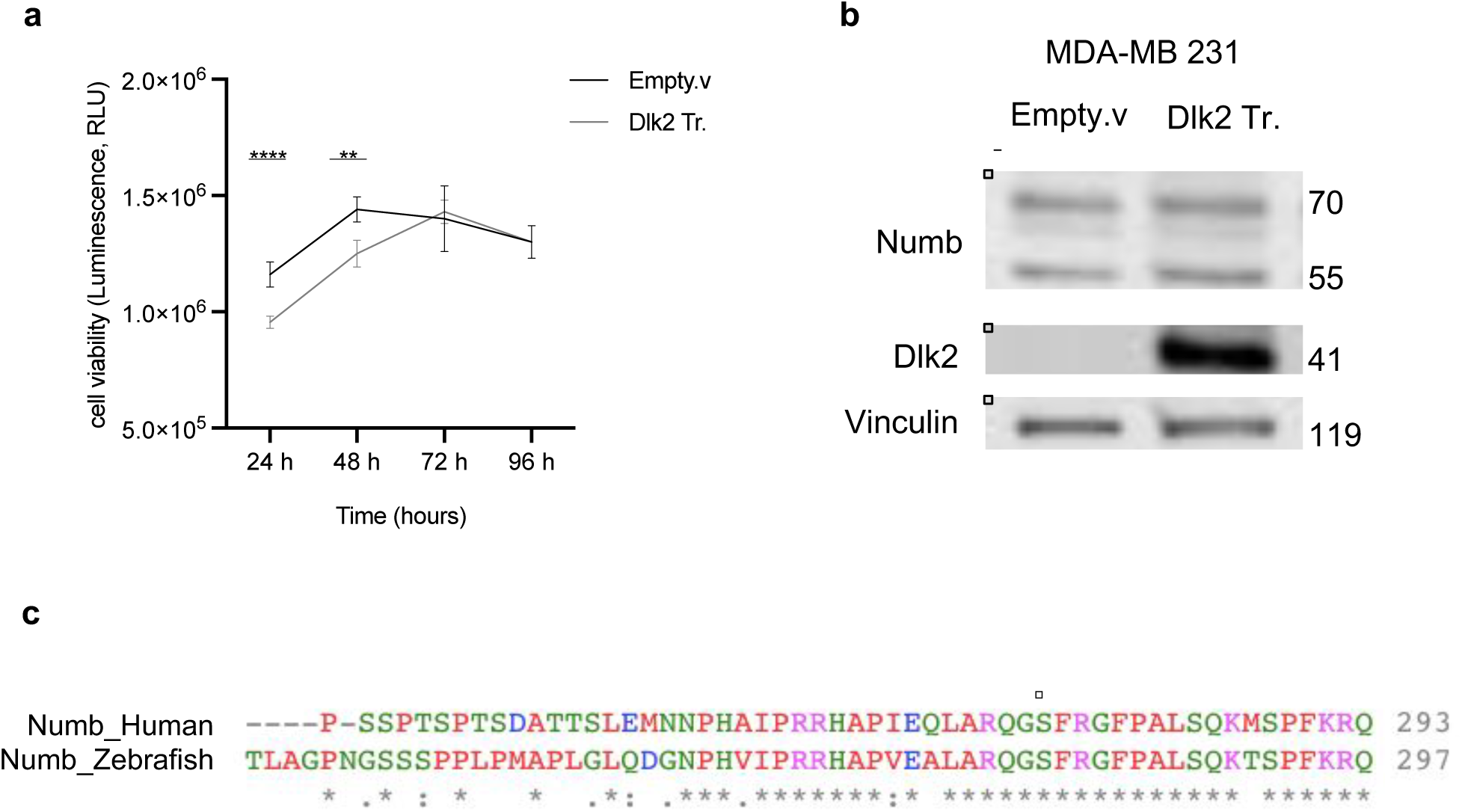
Dlk2 overexpression in human breast cancer cell lines. **a**, Cell viability assay in MCF-7 cells comparing empty vector-transfected control cells (Empty.v) with Dlk2-transfected cells (Dlk2. Tr). Using the CellTiter-Glo assay, luminescence (RLU, relative light unit) was measured as an indicator of cell proliferation. Data are mean ± s.d taken every 24 h in replicate wells (*n* ≥ 3 independent biological replicate experiments). **b,** Overexpression experiment of Dlk2 by transfection in MDA-MB-231 cell line. Western blot of Numb, Dlk2, and Vinculin (loading control). Samples labeled as empty vector (Empty.v) and Dlk2 transfected cells (Dlk2. Tr). (*n* ≥ 3 independent biological replicate experiments). **c,** Protein sequence alignment of Human Numb (P49757) with zebrafish Numb (Q1XD15) using Clustal Omega multiple sequence alignment program. The asterisk symbol indicates positions that have a single, fully conserved residue between the two aligned sequences. The square highlights the alignment with the human Ser276 residue.

